# Evolutionarily related small viral fusogens hijack distinct but modular actin nucleation pathways to drive cell-cell fusion

**DOI:** 10.1101/2020.06.03.130740

**Authors:** Ka Man Carmen Chan, Ashley L. Arthur, Johannes Morstein, Meiyan Jin, Abrar Bhat, Dörte Schlesinger, Sungmin Son, Donté A. Stevens, David G. Drubin, Daniel A. Fletcher

## Abstract

Fusion-associated small transmembrane (FAST) proteins are a diverse family of non-structural viral proteins that, once expressed on the plasma membrane of infected cells, drive fusion with neighboring cells, increasing viral spread and pathogenicity. Unlike viral fusogens with tall ectodomains that pull two membranes together through conformational changes, FAST proteins have short fusogenic ectodomains that cannot bridge the inter-membrane gap between neighboring cells. One orthoreovirus FAST protein, p14, has been shown to hijack the actin cytoskeleton to drive cell-cell fusion, but the actin adaptor-binding motif identified in p14 is not found in any other FAST protein. Here, we report that an evolutionarily divergent FAST protein, p22 from aquareovirus, also hijacks the actin cytoskeleton but does so through different adaptor proteins, Intersectin-1 and Cdc42, that trigger N-WASP-mediated branched actin assembly. We show that despite using different pathways, the cytoplasmic tails of p22 and p14 can be exchanging to create a potent chimeric fusogen, suggesting they are modular and play similar functional roles. When we replace p22’s branched actin nucleator, N-WASP, with the parallel filament nucleator, formin, its ability to drive fusion is maintained, indicating that localized mechanical pressure on the plasma membrane coupled to a membrane-disruptive ectodomain is sufficient to drive cell-cell fusion. This work points to a common biophysical strategy used by FAST proteins to push rather than pull membranes together to drive fusion, one that may be harnessed by other short fusogens responsible for physiological cell-cell fusion.

## Introduction

Aquareovirus and orthoreovirus are two genera of the *Reoviridae* family of segmented double-stranded RNA viruses that form multinucleated syncytium after infection that increases viral spread and pathogenicity (1–4). To drive cell-cell fusion, both aquareovirus and orthoreovirus express a non-structural, fusion-associated small transmembrane (FAST) protein on the plasma membrane of infected cells. The FAST protein is not required for viral entry, and expression of FAST protein alone is sufficient to cause cells to fuse with naïve neighboring cells, forming large multinucleated syncytium (2, 3, 5–12), confirming they are bona-fide cell-cell fusogens. Though they have similar function and topology in the membrane, FAST proteins from aquareovirus and orthoreovirus share minimal sequence identity (13). Based on phylogenetic analysis, they are hypothesized to have evolved from a common, likely non-fusogenic, ancestor 510 million years ago (1, 13, 14). Separate gain-of-function events are believed to have produced fusogenic proteins in both aquareovirus and orthoreovirus, with further divergence or acquisition events resulting in the diversity of FAST proteins found in reoviruses today (13).

Aquareovirus and orthoreovirus FAST proteins are both single pass membrane proteins of fewer than 200 residues comprised of a mostly disordered cytoplasmic tail, a transmembrane domain, and a small ectodomain of fewer than 40 residues (2, 3). The membrane-disruptive ectodomains of FAST proteins typically have solvent-exposed hydrophobic residues and/or myristoylation that are necessary for cell-cell fusion (5, 15–17). In contrast to other cell-cell fusogens that fuse membranes by pulling them together using conformational changes in their ~10 nm tall ectodomains, the minimal ectodomains of FAST proteins have minimal predicted secondary structure, are unlikely to undergo conformational changes to drive membrane fusion (2, 3) and extend only ~1 nm above the bilayer (5, 18). How such short fusogens can overcome the ~2 nm repulsive hydration barrier to reach and fuse with an opposing membrane (5, 18) has been a longstanding question for FAST proteins and other short cell-cell fusogens, such as myomixer and myomaker that are involved in myoblast fusion (19–22).

Recently, we found that the FAST protein from reptilian orthoreovirus, p14, hijacks the host cell actin cytoskeleton to drive cell-cell fusion by forming localized branched actin networks (23). This is accomplished through a c-src phosphorylated tyrosine motif, YVNI, in p14’s disordered cytoplasmic tail that binds to a host adaptor protein, Grb2, which then binds to N-WASP and nucleates branched actin assembly. It is hypothesized that this directly couples local actin-generated forces to push p14’s short, fusogenic ectodomain into the opposing cell’s plasma membrane (23). While all FAST family proteins have similarly short ectodomains, it is unclear if this is a general strategy used by other FAST proteins to drive cell-cell fusion.

Here, we report that a FAST protein from the divergent aquareovirus, p22, also hijacks the host actin cytoskeleton but does so using a molecular strategy distinct from that of orthoreovirus FAST protein p14. Instead of binding to Grb2, we find that p22 binds to Intersectin-1 through an SH3 binding motif in its cytoplasmic tail, which subsequently binds Cdc42 to activate N-WASP-mediated branched actin assembly. We show that despite minimal sequence identity, the p22 cytoplasmic tail can be functionally swapped with that of p14, suggesting that while the cytoplasmic tails of the two FAST proteins evolved independently, they play a similar functional role. By directly coupling the ectodomain with a different actin nucleator, we suggest that actin’s functional role is applying mechanical pressure to a fusogenic ectodomain at the plasma membrane. This biophysical role may be shared across other members of the FAST protein family and could be more generally employed by other cell-cell fusogens.

## Results

### p22 is a fusion-associated small transmembrane protein

To explore whether other FAST proteins might hijack the actin cytoskeleton in the same way as reptilian orthoreovirus FAST protein p14, we first examined the primary sequence of the cytoplasmic tails of FAST proteins from aquareovirus and orthoreovirus, two divergent genera of *Reoviridae*. Surprisingly, the Grb2-binding motif in p14, YVNI, is not found in any other FAST protein (Supp. Figure 1). While other FAST proteins have 1-3 tyrosines in their cytoplasmic tails that could be part of motifs used to bind similar cellular adaptor proteins, p22 has no tyrosines at all (Supp. Figure 1). In fact, p22 from Atlantic Salmon reovirus-Canada 2009, an aquareovirus-A strain, is the most divergent from p14 from reptilian orthoreovirus (13). To investigate if FAST proteins from evolutionarily distant aquareovirus also hijack the actin cytoskeleton to drive cell-cell fusion - and if so, how - we transiently expressed p22 in Vero cells.

Vero cells expressing mCherry-tagged p22 fuse with neighboring cells, forming multinucleated syncytia, consistent with previous reports (Figure 1A and B, Supp. Figure 2A and Video 1) (10). At 36 hours post transfection, multinucleated cells with more than 20 nuclei can be observed (Figure 1C). Similar to other FAST family proteins, p22 is predicted to be a single-pass membrane protein and localize to the plasma membrane in order to drive membrane fusion with neighboring cells. Consistent with that prediction, prior work has shown that p22 localizes to the membrane fraction of lysates (10). Surprisingly, however, when we imaged mCherry-tagged p22 with confocal microscopy, we found that p22 localized minimally to the plasma membrane (Figure 1D) and appeared to reside primarily in intracellular structures (Supp. Figure 2B). By co-expressing with BFP-tagged Rab11 and GFP-tagged EEA1 to label recycling and early endosomes, we found that p22 localizes to endosomes. Similarly, by staining lipid droplets with BODIPY, we found that p22 localizes to the periphery of lipid droplets (Supp. Figure 2C). To determine if p22 is on the plasma membrane, we surface biotinylated p22-GFP expressing cells with cell impermeable NHS-biotin and enriched for biotinylated proteins with streptavidin beads. We found that p22 eluted from streptavidin beads and hence was biotinylated, confirming its presence on the plasma membrane (Figure 1E).

**Figure 1.**
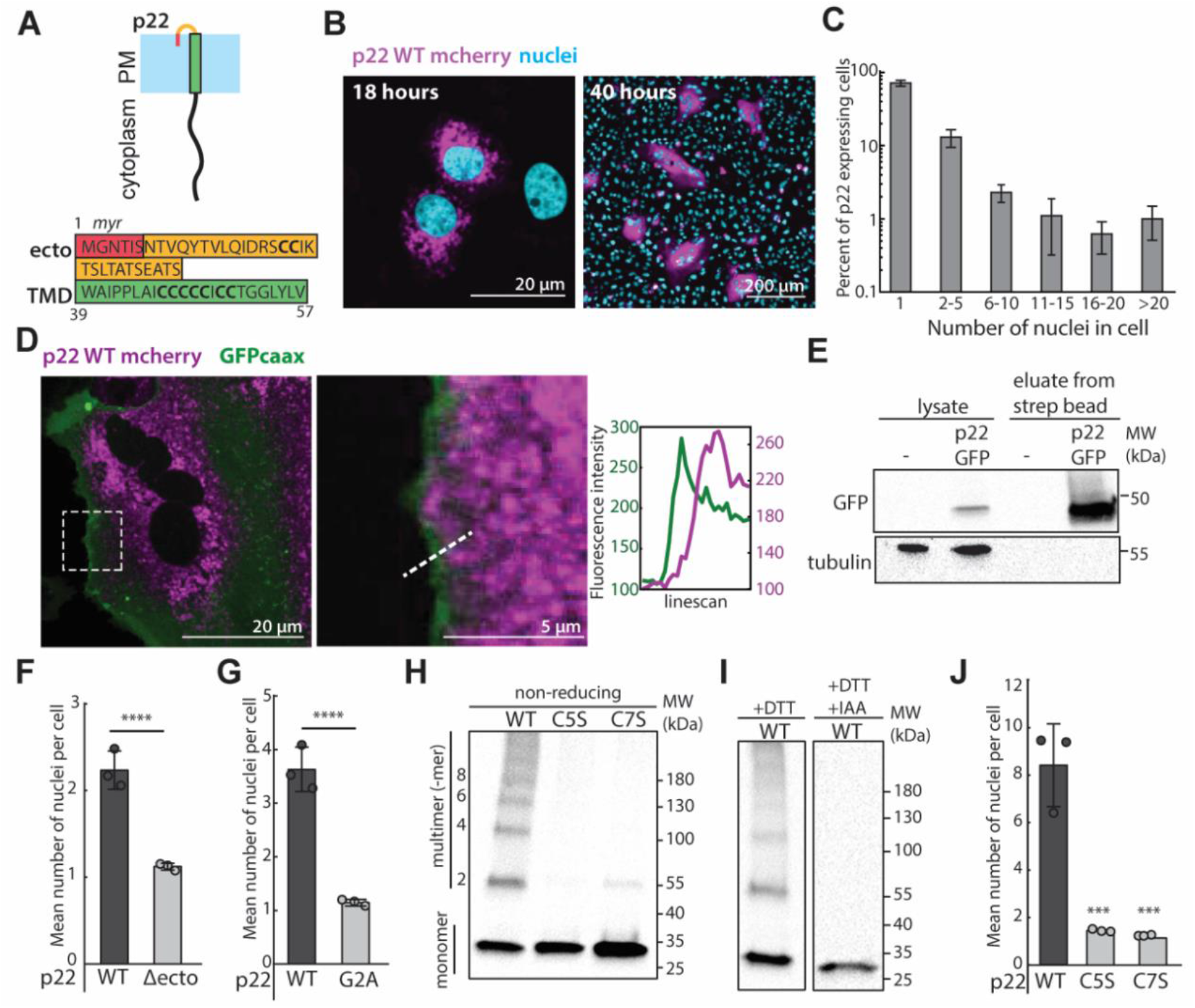
p22 is a membrane protein that multimerizes and drives cell-cell fusion. (A) Diagram of p22 topology on the plasma membrane and amino acid sequence of p22 ectodomain, predicted myristoylation and transmembrane domain. (B) Expression of p22-mcherry (magenta) in Vero cells cause syncytia to form over the ~18-40 hours post transfection. Nuclei are visualized with H2B-GFP (18 hours) and Syto11 (40 hours) (cyan). (C) Nuclei distribution of p22-expressing cells at 36 hours post transfection, error bars indicate average and standard deviation from three independent replicates. (D) Representative confocal images of cells expressing p22-mcherry (magenta) and GFP-caax as a plasma membrane marker (green). Contrast was adjusted in the magnified region. Magnified region and fluorescence intensity of line scan of dotted line is shown. (E) Western blot from surface biotinylation of p22-GFP-expressing cells and non-transfected cells. (F) Mean number of nuclei in p22-WT and p22-Δecto expressing cells from three independent transfections, with error bars representing standard deviations. **** represent p<0.00001 using two-tailed Student’s t-test. (G) Mean number of nuclei in p22-WT and p22-G2A expressing cells from three independent transfections, with error bars representing standard deviations. **** represent p<0.00001 using two-tailed Student’s t-test. (H) Western blot of non-reducing SDS-PAGE gel of myc-tagged p22-WT, p22-TMDC5S, p22-TMDC7S probed with α-myc. (I) Western blot of SDS-PAGE gel of myc-tagged p22-WT, reduced with DTT and capped with iodoacetamide (IAA), and probed with α-myc. (J) Mean number of nuclei in p22-WT, p22-TMDC5S, and p22-TMDC7S expressing cells from three independent transfections, with error bars representing standard deviations. *** represent p<0.0001 using one-way ANOVA with Dunnett’s test.

### p22 has a myristoylated ectodomain and multimerizes with its cysteine-rich transmembrane domain

A common feature of FAST family proteins is a small ectodomain of less than 40 residues with hydrophobic moieties, such as lipidation and/or hydrophobic residues, which are necessary for cell-cell fusion and sufficient to disrupt membranes *in vitro* (2, 3, 16). Similar to other FAST proteins, the p22 ectodomain, which we identified using TMHMM (24), is predicted to be myristoylated according to the Eukaryotic Linear Motif prediction tool (Figure 1A) (25). To determine if myristoylated ectodomain is needed for p22 to drive cell-cell fusion, as it is for p14 and other FAST proteins, we separately truncated the ectodomain (ΔM1-S38, Δecto) and disrupted the myristoylation motif (p22 G2A). While both perturbations did not prevent trafficking to the plasma membrane, both p22 Δecto and p22 G2A were non-fusogenic (Figure 1F and G, and Supp. Figure 2D, E, F, and G). This suggests that the despite its short and unstructured nature, the p22 ectodomain is essential to drive cell-cell fusion and has similar functional motifs as those characterized in FAST proteins from orthoreovirus, including p14 (5, 15, 26).

Surprisingly, the p22 transmembrane domain is predicted, using the same tools as above, to span a cysteine-rich region encoding seven cysteines (Figure 1A) (24). Other FAST family proteins have been shown to multimerize through membrane-proximal and pH-dependent motifs (27, 28), so we hypothesized that these transmembrane cysteines could be used by p22 to multimerize. Using non-reducing SDS-PAGE, we found that p22 WT migrated as a monomeric band at ~27k Da, as a dimer at ~54k Da, and as multimers of dimers at larger molecular weights (Figure 1H). When either the first five cysteines (p22 C5S) or all transmembrane cysteines (p22 C7S) were mutated to serines, higher-order multimerization was abrogated and p22 C5S and C7S migrated primarily as a monomeric band (Figure 1H). Similarly, when the cysteines were reduced with DTT and capped with iodoacetamide, p22 migrated as a single monomeric band (Figure 1I). When multimerization was disrupted in p22 C5S and p22 C7S, trafficking to the plasma membrane was unperturbed, but cell-cell fusion was abrogated (Figure 1J and Supp. Figure 2H and I). Taken together, this data suggests that higher-order multimerization of p22, which could cluster and increase its local concentration on the plasma membrane, is required for cell-cell fusion.

### p22 hijacks the actin cytoskeleton through the ITSN-Cdc42 pathway

To determine if the host cell’s actin cytoskeleton is essential for p22-mediated cell-cell fusion, we broadly inhibited actin polymerization with latrunculin A and found that cell-cell fusion was reduced (Figure 2B and Supp. Figure 3A). When Arp2/3, a key component of branched actin networks, was inhibited with CK-666, p22-mediated cell-cell fusion was also attenuated (Figure 2B and Supp. Figure 3A). However, when formin, nucleators of parallel actin filaments, were inhibited with smiFH2, at more than three times the IC_50_ (29), p22-mediated cell-cell fusion was unchanged (Figure 2B and Supp. Figure 3A). These data suggest that branched actin networks are central to p22-mediated cell-cell fusion.

**Figure 2.**
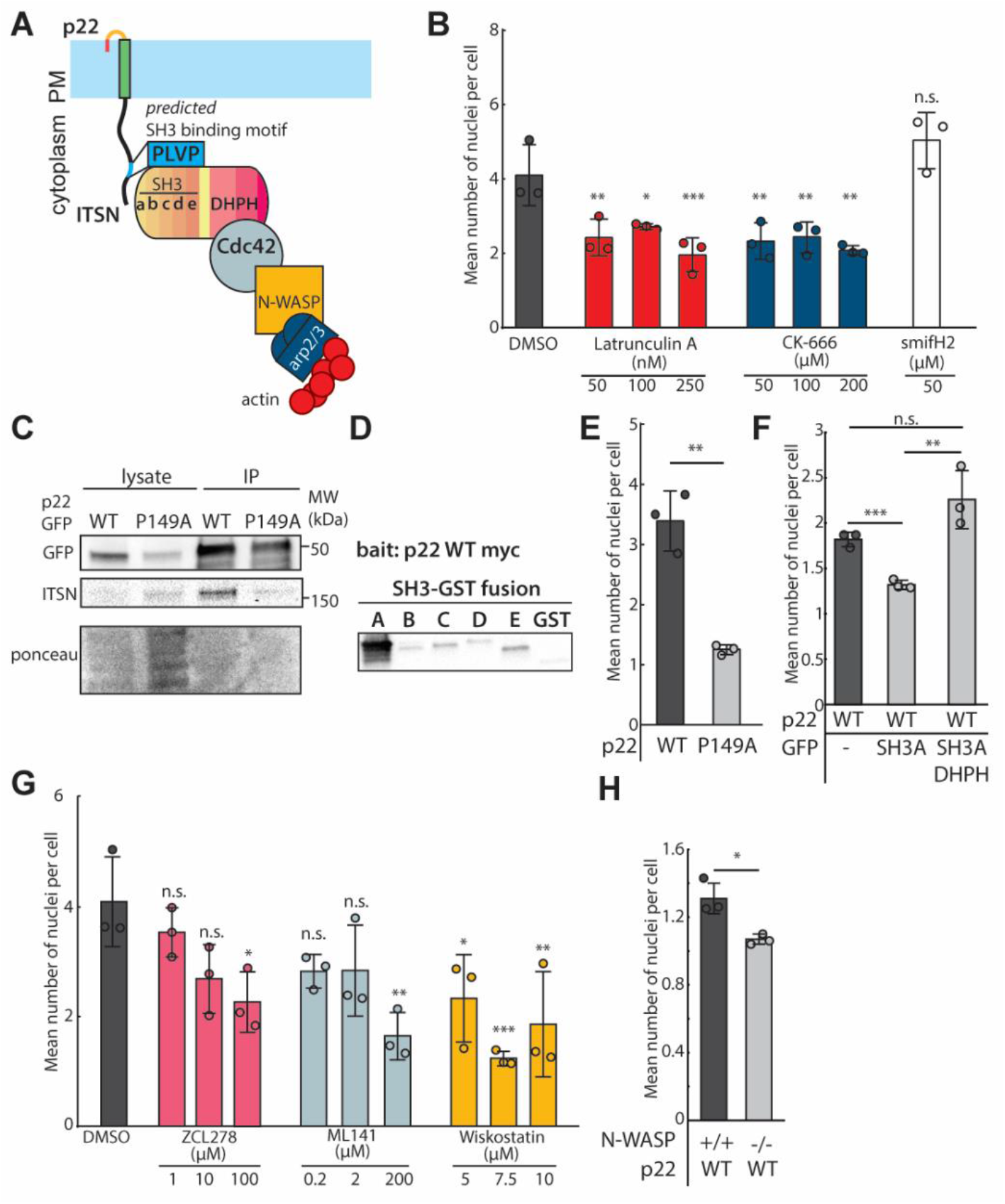
p22 binds to Intersectin-1 to hijack actin assembly and drive cell-cell fusion. (A) Diagram of p22 binding to Intersectin-1 through a SH3 binding motif in its cytoplasmic tail. (B) Mean number of nuclei in p22-WT expressing cells treated with cytoskeletal drugs from three independent transfections, with error bars representing standard deviations. P values are one way ANOVA with Dunnett’s test where n.s. p>0.05, * p<0.05, **p<0.01, *** p<0.001. (C) Western blot of co-immunoprecipitation of GFP-tagged p22-WT and p22-P149A with Intersectin-1. (D) *In vitro* binding of GST-tagged SH3 domains of Intersectin-1 with immunoprecipitated myc-tagged p22-WT. (E) Mean number of nuclei in p22-WT and p22-P149A expressing cells from three independent transfections, with error bars representing standard deviations. ** represent p< 0.01 using two-tailed Student’s t-test. (F) Mean number of nuclei in p22-WT expressing cells co-expressing GFP alone, SH3A-GFP, or SH3A-GFP-DHPH from three independent transfections, with error bars representing standard deviations. ** represent p< 0.01, ***p<0.001, n.s. p>0.05 using two-tailed Student’s t-test. (G) Mean number of nuclei in p22-WT expressing cells treated with drugs from three independent transfections, with error bars representing standard deviations. P values are one way ANOVA with Dunnett’s test where n.s. p>0.05, *p<0.05, **p<0.01, ***p<0.001. (H) Mean number of nuclei N-WASP null mouse embryonic fibroblasts and control cells expressing p22-WT from three independent transfections, with error bars representing standard deviations. P values are two-tailed Student’s t-test where *p<0.01.

Although p22 lacks the phospho-tyrosine motif identified in p14, p22’s disordered cytoplasmic tail might be able to hijack the host cell branched actin cytoskeleton through a different pathway. To investigate this, we first truncated the predicted cytoplasmic tail of p22 (ΔT78-T198, Δcyto) and found that cell-cell fusion was abrogated (Supp. Figure 3B). However, trafficking to the plasma membrane was attenuated (Supp. Figure 2G). To examine how the p22 cytoplasmic tail might couple to branched actin networks to drive cell-cell fusion, we used the Eukaryotic Linear Motif and Scansite 4.0 prediction tools to identify potential binding motifs. We identified an SH3 domain binding motif (PXXP) in the p22 cytoplasmic tail (P146-P149, PLVP) that was predicted to bind to Intersectin-1 (Figure 2A). Intersectin-1 is an endocytic scaffolding protein that consists of two EH domains that bind to endocytic components, DH and PH domains, and five SH3 domains that couple to adaptor proteins. Any or all of these five SH3 domains could bind to p22’s SH3 binding motif and may be important for p22-mediated cell-cell fusion.

Using co-immunoprecipitation to verify this prediction, we found that Intersectin-1 does indeed directly bind to p22 WT (Figure 2C). To determine which of the five SH3 domains p22 binds to, we purified each SH3 domain as a GST-fusion protein and incubated it with immunoprecipitated, myc-tagged p22 (Supp. Figure 3C). We found that p22 strongly binds to SH3 domain A and weakly to the other four SH3 domains of Intersectin-1 (Figure 2D). To determine if p22 WT co-localizes with Intersectin-1 in live cells, we transfected genome-edited cells expressing endogenously mEGFP-tagged Intersectin-1 genome-edited cells with HaloTag-tagged p22-WT. Because p22 WT is undetectable on the plasma membrane via confocal microscopy, we imaged endosomes and the periphery of lipid droplets where p22 is primarily localized and found that Intersectin-1 co-localizes with it (Supp. Figure 3D). When the SH3 domain binding motif is disrupted (P149A), p22 P149A is still present in these Intersectin-1 endosomes and lipid droplets and it is still trafficked to the plasma membrane as evidenced by surface biotinylation (Supp. Figure 3D and E). However, p22 P149A no longer directly binds to Intersectin-1 (Figure 2C), and cell-cell fusion is abrogated (Figure 2E, Supp. Figure 3F), suggesting it plays a key role in cell-cell fusion. To confirm that direct coupling of p22 to Intersectin-1 is needed for p22-mediated cell-cell fusion, we over-expressed the SH3 domain A of Intersectin-1 to compete with endogenous Intersectin-1 and found that the extent of p22-mediated cell-cell fusion was reduced (Figure 2F and Supp. Figure 3G). Taken together, these findings indicate that p22 binding to Intersectin-1 is necessary to drive cell-cell fusion.

The DH and PH domains of Intersectin-1 act as a guanine-exchange-factor (GEF) for the small GTPase, Cdc42, which activates N-WASP to nucleate branched actin network assembly. This GEF activity has been implicated in actin comet tail formation in Vaccinia viruses and is sufficient to trigger assembly of actin comet tails from purified proteins *in vitro* (30–32). To determine if these downstream effectors are necessary for p22-mediated cell-cell fusion, we specifically targeted the Intersectin-Cdc42 binding site with the inhibitor ZCL278 (33) and found that cell-cell fusion was impaired (Figure 2G and Supp. Figure 3H). To determine if GEF activity is sufficient to drive p22-mediated cell-cell fusion, we over-expressed a fusion protein consisting of only the SH3 domain A, the DH and PH domains of Intersectin-1. This restored p22-mediated cell-cell fusion (Figure 2F and Supp. Figure 3G). We next inhibited Cdc42 GTPase activity allosterically with ML141 and found that it also blocked p22-mediated cell-cell fusion (Figure 2H and Supp. Figure 3H). When N-WASP was inhibited with Wiskostatin, p22-mediated cell-cell fusion was also inhibited (Figure 2H and Supp. Figure 3H). To confirm that N-WASP was specifically required for nucleation of branched actin networks by p22, we expressed p22 in N-WASP-null mouse embryonic fibroblasts (MEFs). Despite the lower transfection efficiency in MEFs, we found that p22-mediated cell-cell fusion was attenuated in N-WASP-null MEFs (Figure 2H and Supp. Figure 3I). Altogether, these data show that p22, like p14, hijacks branched actin network assembly to drive cell-cell fusion but uses a distinct pathway. Instead of using a phosphorylation-dependent motif to bind to adaptor proteins, p22 binds to Intersectin-1 through an SH3 binding motif and then relies on Intersectin-1 GEF activity to activate Cdc42 activity to locally trigger N-WASP-mediated actin assembly.

### p14 and p22 are modular cell-cell fusogens

The functional similarities of the molecular pathways used by p22 and p14 suggest that the cytoplasmic tails of FAST proteins from aquareovirus and orthoreovirus might be modular components of a minimal cell-cell fusogen. To directly test if p22 and p14 are modular, we swapped p22’s cytoplasmic tail with that of p14 to create a chimeric fusogen that is comprised of the p14 ectodomain, p14 transmembrane domain, and p22 cytoplasmic tail (p14/p22 chimera, Figure 3A). Since trafficking to the plasma membrane for Type III integral membrane proteins is primarily determined by their transmembrane domain, we hypothesized that the p14/p22 chimera, with p14’s transmembrane domain that localizes readily to the plasma membrane (Figure 3B), will be expressed at higher levels on the plasma membrane. Consistent with this, the p14/p22 chimera showed higher plasma membrane localization than wild-type p22 (Figure 3B and C). 24 hours post transfection, we found that the p14/p22 chimera expressing cells have significantly more nuclei and fuse more readily than those expressing wild-type p14 or p22 (Figure 3D), forming large syncytia with more than 80 nuclei (Figure 3E, and Supp. Figure 4A). Due to extensive fusion, merging of transfected cells with non-transfected cells diluted the p14/p22 chimera and decreased its concentration (Figure 3F, and Supp. Figure 4B). Despite the lower surface density, the p14/p22 chimera is so potent that we had to assay cell-cell fusion 12 hours earlier than in our standard assay in order to be able to quantify the number of nuclei in p14/p22 chimera-expressing cells.

**Figure 3.**
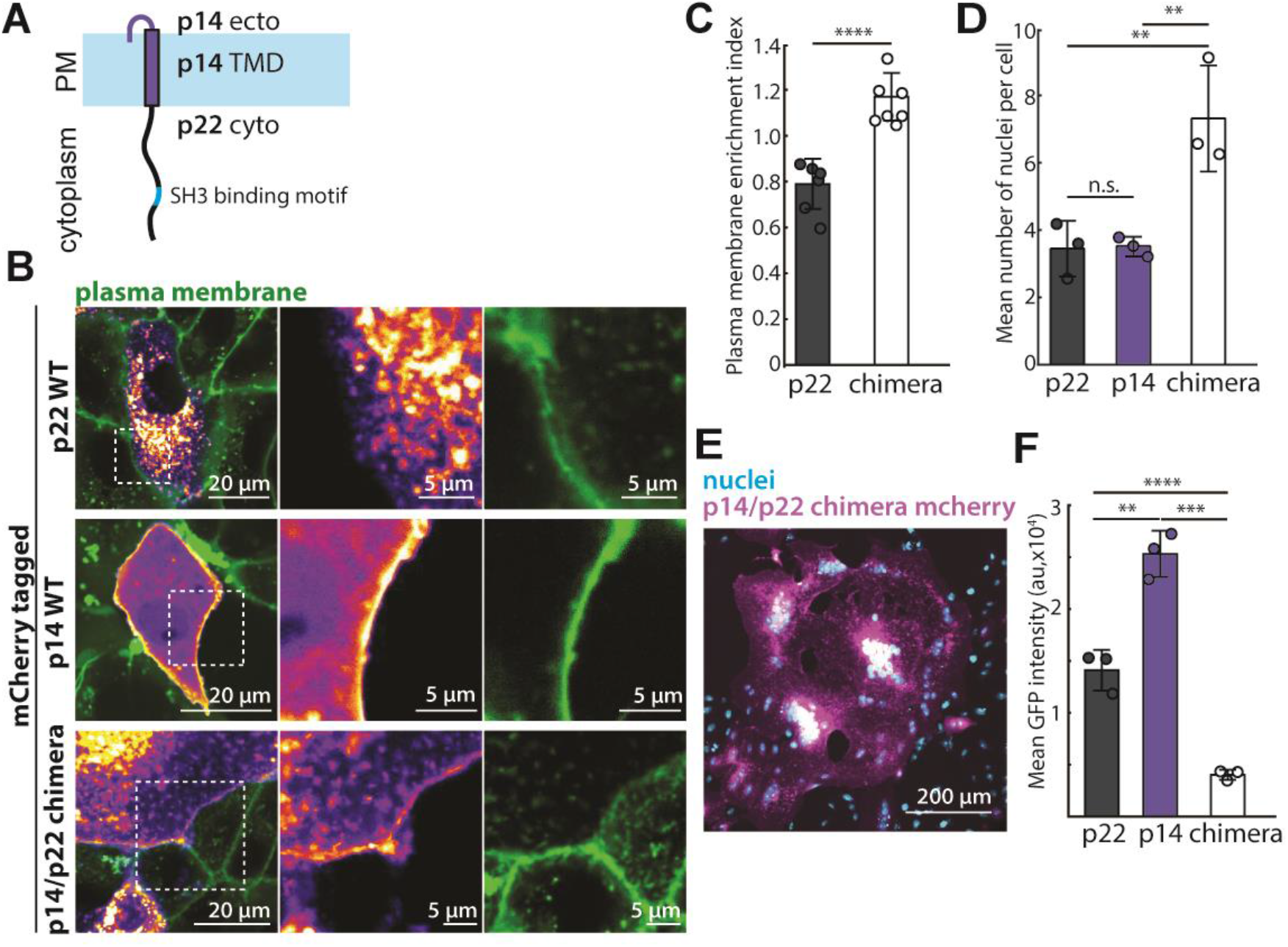
p14 and p22 are modular cell-cell fusogens and their cytoplasmic tails can be swapped. (A) Diagram of chimeric fusogen with p14 ectodomain, p14 transmembrane domain and p22 cytoplasmic tail. (B) Confocal images of mCherry tagged p22-WT, p14-WT and p14/p22 chimera with plasma membrane labeled with CellMaskDeepRed (green). Boxed regions are magnified. (C) Average plasma membrane enrichment index of mCherry tagged p22-WT and p14/p22 chimera from three independent transfections and error bars represent standard deviation 24 hours post transfection. P values are two-tailed Student’s t-test where ****p<0.0001. (D) Mean number of nuclei in p22-WT, p14-WT and p14/p22 chimera expressing cells 24 hours post transfection from three independent transfections, with error bars representing standard deviations. P values are one-way ANOVA with Tukey’s test where n.s. p>0.05, **p<0.01. (E) Representative confocal image of p14/p22 chimera mCherry (magenta) cell with nuclei labeled with Hoechst 33342 (cyan) at 24 hours post transfection. (F) Mean GFP intensity in each cell expressing GFP-tagged p22-WT, p14-WT and p14/p22 chimera 24 hours post transfection from three independent transfection and error bars represent standard deviations. P values are two-tailed Student’s t-test where **p<0.01, ***p<0.001, ****p<0.0001.

Why is the p14/p22 chimera more fusogenic than either wild-type p14 or p22? Based on previous work, we know that p14 drives cell-cell fusion by binding to Grb2 through a motif in its cytoplasmic tail that needs to be phosphorylated by c-src at the plasma membrane (23) and p14’s fusogenicity is attenuated when this binding motif is dephosphorylated. Furthermore, the cytoplasmic tails of a fraction of p14 are cleaved, rendering it non-fusogenic (23). In contrast, p22 is minimally cleaved (Figure 3B) and binds to its adaptor protein, Intersectin-1, in a constitutively active manner. However, its fusogenicity is likely restricted by limited trafficking to the plasma membrane, which might be limited by the non-conventional, cysteine-rich transmembrane domain (Figure 1D). Therefore, when the p22 cytoplasmic tail is ligated with p14’s fusogenic ectodomain and transmembrane domain, the p14/p22 chimera is readily trafficked to the plasma membrane with constitutively active coupling to actin assembly, resulting in a more potent fusogen than either individually.

### Replacing the p14/p22 chimera’s branched actin nucleator with formin preserves cell-cell fusion

The fusogenicity of the p14/p22 chimera suggests that FAST proteins of both aquareovirus and orthoreovirus are modular cell-cell fusogens and their specific molecular identity is not crucial to drive cell-cell fusion. Their cytoplasmic tails use different molecular strategies to accomplish the same task of coupling actin assembly with a membrane-disruptive ectodomain. If the primary role of actin assembly is to apply localized pressure at the plasma membrane and “push” the membrane-disruptive ectodomain into the opposing plasma membrane, could other force-generating actin nucleators also drive cell-cell fusion? Branched actin networks can exert up to 5 nN or about 1250 nN/μm^2^ as they grow, a filopodium can exert up to only 10 pN (34–37). However, the diameter of a filopodium is 100-300 nm and hence can exert localized pressures ranging from ~140-1270 nN/μm^2^ (37–40), comparable to branched networks in terms of local pressure.

To determine if the pressure generated by formin-based actin nucleation is sufficient to drive cell-cell fusion, we first rendered p14/p22 chimera non-fusogenic by disrupting the SH3 binding motif. We then coupled the non-fusogenic mutant of p14/p22 chimera to a constitutively active formin (ΔGBD-mDia2), which is involved in filopodia formation, using FKBP-FRB tags (Figure 4A). Co-expression of p14/p22 chimera-P149A-FKBP with an FRB-ΔGBD-mDia2 led to filopodia formation (Figure 4B), consistent with previous reports expressing ΔGBD-mDia2 (41). Upon addition of the rapamycin analog, the p14/p22 chimera-P149A-FKBP clustered and enriched at the tip of the filopodia (Figure 4B and C, Video 2 and 3) and the filopodia elongated (Video 2 and 3). The enrichment and clustering of p14/p22 chimera-P149A-FKBP at the filopodia tips is likely due to coupling to the FRB-tagged formins that sit on the ends of actin filaments where they drive filament polymerization and subsequent filament bundling (42–45). Remarkably, this coupling of ΔGBD-mDia2 with the p14/p22 chimera-P149A-FKBP is sufficient to drive cell-cell fusion (Figure 4D and Supp. Figure 5). Overall, this suggests that force generated by the local assembly of a different actin structure is sufficient to drive cell-cell fusion, supporting the hypothesis that FAST proteins are minimal fusogens that couple membrane-disruptive ectodomains with localized pressure exerted on the plasma membrane.

**Figure 4.**
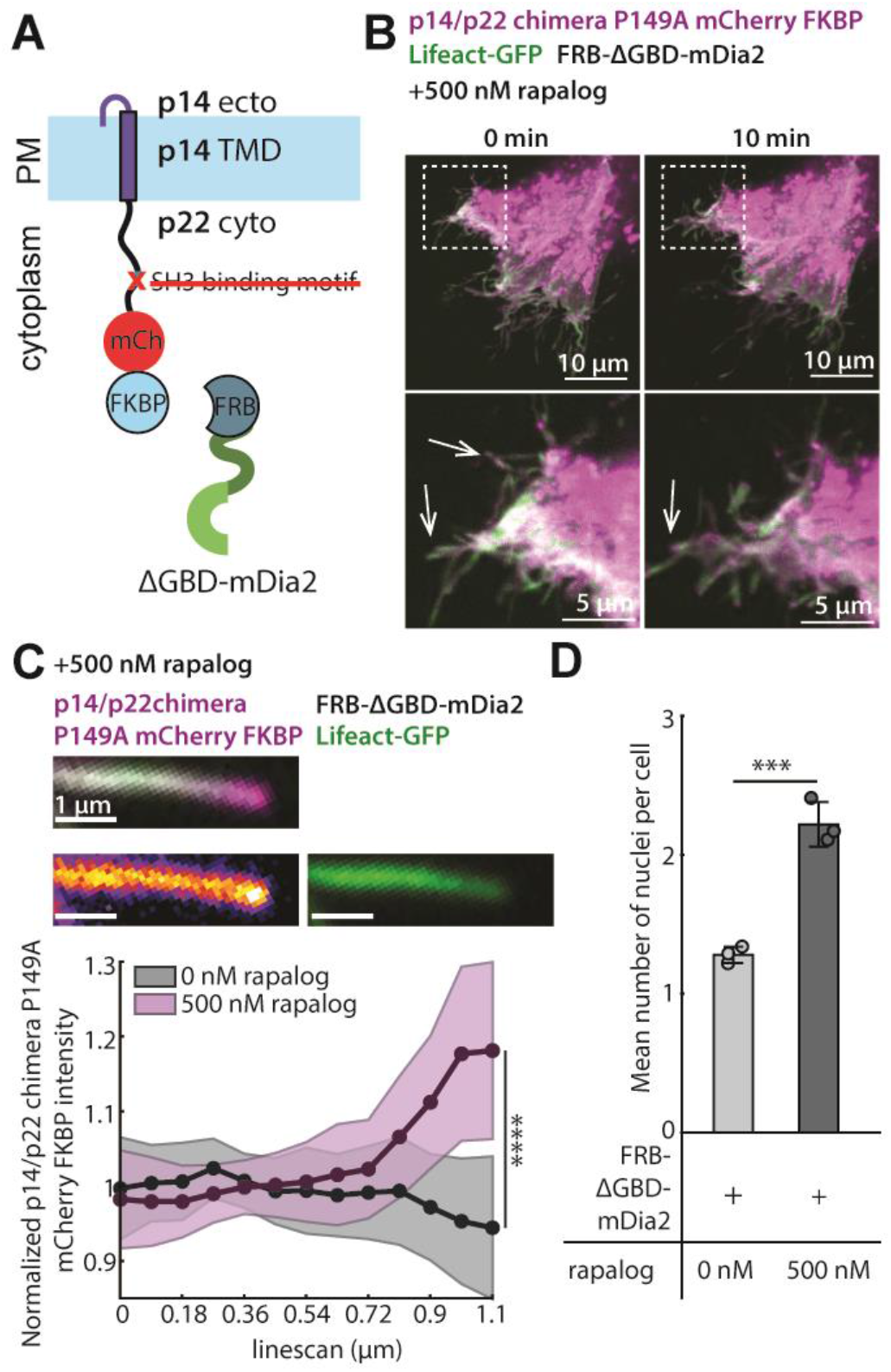
Replacing the p14/p22 branched actin nucleator with a formin is sufficient to drive cell-cell fusion. (A) Diagram of p14/p22 chimera with FKBP at C-terminus and P149A mutation with FRB-tagged ΔGBD-mDia2. (B) Confocal images of p14/p22 chimera P149A FKBP-mCherry (magenta) co-expressed with FRB-ΔGBD-mDia2 and Lifeact-GFP (green) cell at 0 min and 10 min after addition of 500 nM of rapalog. Filopodia-like protrusions before and after rapalog addition is denoted with arrows. Box regions magnified. (C) Representative confocal merged image of a filopodia-like protrusion from cell expressing p14/p22 chimera-P149A-FKBP-mCherry (magenta), Lifeact-GFP (green), and FRB-ΔGBD-mDia2 10 minutes after addition of 500 nM rapalog. Each channel is shown with p14/p22 chimera-P149A-FKBP-mCherry (fire) and Lifeact-GFP (green). Average normalized fluorescence intensity of p14/p22 chimera-P149A-FKBP-mCherry along the length before (n=33 filopodia-like protrusions) and 10 min after (n=36 filopodia-like protrusions) addition of 500 nM rapalog. Standard deviation above and below the average are shown. **** p<0.0001 by Student’s t-test on the last datapoint. (D) Average mean number of nuclei in p14/p22 chimera-P149A-FKBP and FRB-ΔGBD-mDia2 expressing cells with and without rapalog from three independent transfections, with error bars representing standard deviations. P values are two-tailed two-sample Student’s t-test where ***p<0.001.

## Discussion

FAST proteins are a unique family of fusogens that lack the large ectodomains and ability to undergo conformational changes of better-studied fusogens, such as hemagglutinin and the cell-cell fusogens, Eff-1 and Hap2 (46–51). We previously found that one member of this family, p14, nucleates branched actin networks assembly by binding the adaptor protein Grb2 to a phospho-tyrosine motif in its cytoplasmic tail, which subsequently binds the nucleation-promoting factor N-WASP (23). While the FAST family proteins are thought to have evolved from a common ancestor over 500 million years ago and now show little sequence similarity, we hypothesized that other fusogens in the FAST family also harness the actin cytoskeleton to push their similarly short ectodomains into the opposing bilayer. However, analysis of the cytoplasmic domains of all known FAST proteins revealed that none had the required Grb2-binding sequence. This led us to investigate how an evolutionary distant FAST protein from aquareovirus, p22, is able to drive cell-cell fusion without using the molecular strategy employed by p14.

In this study, we found that p22 uses a distinct phosphorylation-independent SH3 binding motif to bind to Intersectin-1 and subsequently Cdc42 to hijack N-WASP mediated actin assembly for cell-cell fusion. Interestingly, this SH3 motif is conserved in the FAST protein of Turbot reovirus, another aquareovirus-A strain (NS22, Supp. Figure 6). However, such an Intersectin-1 binding motif is not found in other orthoreovirus and aquareovirus species, which highlights the divergence of FAST proteins and raises the possibility that the other FAST proteins (p16, p10, p13, p15) may hijack different, yet unidentified host factors to drive cell-cell fusion.

Although neither the Grb2 nor Intersectin-1 binding motifs are conserved beyond closely-related aquareovirus and orthoreovirus, FAST proteins appear to be modular cell-cell fusogens with a common functional role for their cytoplasmic tails. Both p22 and p14 cytoplasmic tails couple to N-WASP-nucleated branched actin assembly and can be swapped to create a functional p14/p22 chimeric fusogen. Interestingly, the p14/p22 chimera reported here is substantially more potent than wild-type p14 or p22. This suggests that the fusogenicity of FAST proteins that perhaps selective pressures limit the potency of FAST protein fusion during viral infection in an organism (4). Although the actin cytoskeleton has yet to be implicated in other FAST proteins, similar chimeric fusogens with p10 from avian reovirus and p15 from baboon reovirus have also been shown to be functional (2, 27), further supporting the idea that FAST family fusogen domains are modular.

The modularity of FAST family proteins indicates that the specific identity of host molecular players hijacked by the fusogens is secondary to their biophysical roles. Both p14 and p22 hijack N-WASP, which nucleates branched actin networks that can exert pressures up to 1250 nN/ μm^2^. However, when we replace N-WASP with a formin, which nucleates parallel actin bundles instead of branched actin networks, we find that cell-cell fusion is preserved. While branched actin structures have been implicated during other physiological and pathological cell-cell fusion processes, this is the first description (albeit synthetic) of filopodia-mediated cell-cell fusion.

Overall, our findings support a model of cell-cell fusion in which the minimal requirements for fusion are mechanical pressure to bring two plasma membranes together, regardless of how the force is generated, coupled to a membrane-disruptive ectodomain. Localized assembly of actin beneath a membrane-disruptive ectodomain could be a fundamental strategy for overcoming the fusion energy barrier, one that is used not only by FAST family proteins but also by other cell-cell fusogens that do not rely on conformational changes to pull membranes together, such as myomaker and myomixer.

## Materials and Methods

### Molecular Cloning

Aquareovirus fusion associated small transmembrane protein, p22 (Atlantic salmon reovirus Canada-2009, Accession number: C0L0N0), was synthesized and inserted into mammalian expression vector pcDNA3.1 with C-terminus tags (mcherry, eGFP, myc). Point mutations and truncations were introduced with primers and verified with Sanger sequencing. pcDNA3.1-p14-mcherry was used as previously described. Chimera with p14 ectodomain and transmembrane domain (1-57 AA) and p22 cytoplasmic tail (58-198 AA) was synthesized with Gibson assembly with C-terminus tags (mcherry, eGFP, myc).

GFP-EEA1 WT was a gift from Silvia Corvera (Addgene plasmid # 42307).

GFP-Intersectin Long, Intersectin-1 I SH3 A domain (human), Intersectin-1 I SH3 B domain (human), Intersectin-1 I SH3 C domain (human), Intersectin-1 I SH3 D domain (human), and Intersectin-1 I SH3 E domain (human), were gifts from Peter McPherson (Addgene plasmids #47395, #47413, #47414, #47415, #47416, #47417).

ΔGBD-mDia2 (258-1171 AA) was amplified from pCMV-eGFP-mDia2 (a kind gift from Scott Hansen) and inserted with a N-terminus FRB tag into pcDNA3.1.

SH3 A domain from human Intersectin-1 (740-816 AA) was amplified and inserted with a C-terminus eGFP tag into pcDNA3.1. DH and PH domains from human Intersectin-1 (1226-1573 AA) was amplified and inserted downstream of SH3 A domain and eGFP into pcDNA3.1.

Rab11a (human) was synthesized as a gblock from IDT and inserted with N-terminus mTagBFP2 tag.

### Cell culture, transfection and generation of mutant SH3 overexpression cells

Vero cells were obtained from UC-Berkeley Cell Culture Facility. Vero cells were grown in DMEM (Life Technologies) supplemented with 10% heat-inactivated FBS (Life Technologies), 10% non-essential amino acids (Life Technologies), and 1% Pen-Strep (Life Technologies), at 37 °C, 5% CO2. N-WASP −/− and +/+ mouse embryonic fibroblasts cells were a kind gift from Scott Snapper. and were grown in DMEM (Life Technologies) supplemented with 10% heat-inactivated FBS (Life Technologies), and 1% Pen-Strep (Life Technologies), at 37 °C, 5% CO2.

Cells were negative for mycoplasma as verified with Mycoalert mycoplasma detection kit (Lonza).

Cells were transfected with FuGENE HD (Promega) according to manufacturer’s instructions.

### Genome-edited human induced pluripotent stem cells (hiPSCs)

WTC10 hiPSC line was obtained from the Bruce Conklin Lab. hiPSCs were cultured on Matrigel (hESC-Qualified Matrix, Corning) in StemFlex medium (Thermo Fisher) with Penicillin/Streptomycin in 37°C, 5% CO2. Cultures were passaged with Gentle Cell Dissociation reagent (StemCell Technologies) twice every week.

The AP2M1 gene was edited in WTC10 hiPSCs as previously described using TALENs targeting exon 7 of AP2M1 gene (52). Both alleles of AP2M1 were tagged with tagRFP-T. Cas9-crRNA-tracrRNA complex electroporation method was used to edit ITSN1 gene in AP2M1-tagRFP-T genome edited hiPSCs. S. pyogenes NLS-Cas9 was purified in the University of California Berkeley QB3 MacroLab. TracrRNA and crRNA targeting AGGTGTTGGAAACTGAGCCA sequence in the immediate vicinity of the start codon of ITSN1 were purchased from IDT. Gibson assembly (New England Biolabs) was used to construct a donor plasmid containing the pCR8/GW/TOPO (Thermo Fisher) plasmid backbone, the mEGFP gene followed by a AAGTCCGGAGGTACTCAGATCTCGAGG linker sequence, and 450/447 base pair homology arm sequences. Three days after electroporation (Lonza, Cat#: VPH-5012) of Cas9-crRNA-tracrRNA complex and donor plasmid, the GFP positive cells were single cell sorted with a BD Bioscience Influx sorter (BD Bioscience) into Matrigel-coated 96-well plates. Clones were confirmed by PCR and sequencing of the genomic DNA locus around the mEGFP insertion site. Both alleles of ITSN1 were tagged with mEGFP in the hiPSC line used in this study.

### hiPSCs sample preparation

2 days before fixation, hiPSCs were seeded on Matrigel-coated 4-well chambered cover glass (Cellvis). 20 hours before fixation, the p22 expressing plasmids were transfected using Lipofectamin Stem Transfection Reagent (Thermo Fisher). Halotag was labeled by JF635-HaloTag ligand (53). Cells were incubated in StemFlex medium with 100 mM JF635-HaloTag for 45min and the unbound ligands were washed away by three times of 5 min incubation in prewarmed StemFlex medium. Cells were fixed by 4% PFA in DPBS (Gibco) for 20 min and washed by DPBS three times.

### Imaging

All live cells were maintained at 37°C, 5% CO_2_ with a stage top incubator (okolab) during imaging.

For confocal microscopy, cells were imaged with a spinning disk confocal microscope (Eclipse Ti, Nikon) with a spinning disk (Yokogawa CSU-X, Andor), CMOS camera (Zyla, Andor), and either a 4x objective (Plano Apo, 0.2NA, Nikon), 20x objective (Plano Fluor, 0.45NA, Nikon), 40x objective (Plano Fluor, 0.6NA, Nikon) or a 60x objective (Apo TIRF, 1.49NA, oil, Nikon). For total internal reflection fluorescence (TIRF) microscopy, cells were imaged with TIRF microscope (Eclipse Ti, Nikon), 60x objective (Apo TIRF, 1.49NA, oil, Nikon) and EMCCD camera (iXON Ultra, Andor). Both microscopes were controlled with Micro-Manager. Images were analyzed and prepared using ImageJ (National Institutes of Health).

### Nuclei count

26,700 Vero cells were transfected with specified plasmids with FuGENE HD (Promega) and immediately plated onto fibronectin coated glass-bottomed dishes. 82,600 N-WASP −/− and +/+ mouse embryonic fibroblasts were transfected with p22 WT with Lipofectamine 3000 and immediately plated onto fibronectin coated glass bottomed dishes. For Vero cells expressing p22 WT and point mutations of p22 WT, cells were fixed 36 hours post transfection and imaged, while for p14/p22 chimera and N-WASP −/− and +/+, cells were fixed 22-24 hours post transfection. Vero cells co-expressing p14/p22 chimera-P149A-FKBP with either FRB-ΔGBD-mDia2 were transfected with FuGene HD. 12-14 hours post transfection, 500 nM of rapalog (AP21967, Takara Bio) was added to media. Cells were fixed 24 hours post transfection. Twenty-five to thirty fields of view from each of three independent transfections were collected.

Micrographs were processed in FIJI and cytoplasmic p22 fluorescence intensity was used to make a binary mask of cell bodies. Nuclei masks were made from cells stained with 5 *μ*M Syto11 (Thermo Fisher), Hoechst 33342 (Thermo Fisher), or from the inverted mask of cytoplasm fluorescence. Nuclei per cell expressing fluorescently-tagged constructs were counted either manually or using the Speckle Inspector FIJI plugin (http://www.biovoxxel.de/). Cells expressing the p14/p22 chimera were masked using pixel classification in Ilastik (https://www.ilastik.org/) and nuclei were counted as above. Cells with more than 2 nuclei were considered multinucleated.

### Plasma membrane enrichment quantification

The plasma membrane of p22-mCherry and p14/p22-Chimera-mCherry expressing cells were labeled with CellMaskDeepRed (ThermoFisher). Cells were imaged with spinning disk confocal microscopy with a 60x objective. A linescan spanning the plasma membrane and 600-1000 nm proximal to the plasma membrane was analyzed in ImageJ. The plasma membrane location is determined by the maximum fluorescence intensity of CellMaskDeepRed, and the plasma membrane enrichment index is defined as the fluorescence intensity of mCherry-tagged proteins at the plasma membrane, normalized to the average fluorescence intensity of the protein in the cytosol.

### Filopodia enrichment quantification

Vero cells co-expressing p14/p22 chimera-P149A-FKBP, FRB-ΔGBD-mDia2 and Lifeact-GFP were imaged with spinning disk confocal microscopy with a 60x objective at 37°C, 5% CO_2_. 500 nM rapalog (AP21967, Takara Bio) was added to the media, and cells were imaged for 15 min. Linescans spanning 1.1 μm from the tip of filopodia were analyzed using ImageJ. The last 270 nm of filopodia is defined as the tip of the filopodia and the remaining 900 nm is defined as the length of the filopodia. Fluorescence intensity of p14/p22 chimera-P149A-mCherry-FKBP along the length of the filopodia is normalized to the average fluorescence intensity of the entire length of the filopodia. Approximately 30 filopodia at 0 min and 30 filopodia at 10 min after addition of rapalog, all selected at random, were analyzed.

### Surface biotinylation

Cells were grown in 100 mm dish or T25 flask to 50% confluency and transfected using FuGENE reagent. After 24 hours cells were washed with PBS (pH = 8) three times on ice and incubated with 1 mg NHS-Biotin in 0.5 mL PBS (pH = 8) for 1 hour on ice. The biotinylation reaction was quenched with 100 mM glycine in PBS for 10 min at room temperature. Cells were washed three times with 100 mM glycine in PBS and lysed with 1 mL lysis buffer (1x RIPA buffer supplemented with 1x HALT protease inhibitor (Thermo Fisher Scientific)) for 30 minutes on ice. Cells were scraped into 1.5 mL microcentrifuge tube and sonicated on ice for 3 minutes. Cell debris was pelleted for 5 min at 17,900 rcf at 4°C. 30 μl of streptavidin magnetic beads (ThermoFisher) were washed with RIPA buffer at 4°C, the supernatant was added and incubated overnight. Beads were washed 5 times with RIPA buffer, suspended in Laemmli sample buffer (30 ul), denatured at 95°C for 5 min and separated on 4-20% acrylamide gradient gels by SDS-PAGE. Proteins were transferred onto nitrocellulose membrane with iBlot (Thermo Fisher Scientific). The membrane was blocked with 5% milk powder in TBST for 1 hour and probed with primary antibodies α-GFP (1:5000, g1544, Sigma Aldrich), α-myc (1:5000, 9e10, Sigma Aldrich), α-tubulin (1:5000, Clone YL1/2, Thermo) in 5% milk in TBST overnight at 4°C. The membrane was then probed with secondary antibodies, α-rabbit HRP (1:5000, 65-6120, Thermo Fisher), α-mouse HRP (1:5000, Jackson Labs), α-rat AlexaFluor 647 (1:5000, Life Technologies). Western blots were imaged on ChemiDoc (Bio-Rad).

### Non-reducing SDS-PAGE, reduction and alkylation

Vero cells were transfected with p22 WT, C5S and C7S with myc tag. Vero cells were harvested 24 hours post transfection. Cells were washed with PBS, and lysed with lysis buffer (150 mM NaCl, 25 mM HEPES pH 7.4, 1 mM EDTA, 0.5% NP-40, 1x HALT protease inhibitor (Thermo Fisher Scientific) for 30 min at 4°C, and scraped and bath sonicated in ice for 3 min. Cell debris was pelleted at 17,900 rcf for 10 min. To enrich for p22, supernatant was incubated with 5 μl of myc-Trap beads (Chromotek) overnight at 4°C. The beads were washed with lysis buffer five times. For p22 WT, C5S and C7S to be analyzed with non-reducing SDS-PAGE, samples were boiled in Laemmli sample buffer without reducing agents and separated on 4-20% acrylamide gradient gels by SDS-PAGE. For p22 WT reduced with DTT, samples were boiled in Laemmli sample buffer with 350 mM DTT. To cap cysteines in p22 WT, myc-Trap beads (Chromotek) were washed three times in alkylation buffer, 100 mM Tris, pH 8.0, 50 mM DTT before incubating at 85°C for 10 min. Reduced cysteines were capped immediately with 100 mM iodoacetamide (Sigma) for 15 min at room temperature in the dark. The eluted protein was boiled in Laemmli sample buffer and separated on 4-20% acrylamide gradient gels by SDS-PAGE. Proteins were transferred onto nitrocellulose membrane 16 hours at 4°C at 40 mAmps. The membrane was blocked 5% milk in TBST for 1 hour and probed with primary antibody, α-myc (1:2500, 9E10, Sigma) in 5% milk in TBST overnight at 4°C. The membrane was then probed with secondary antibody, α-mouse HRP (1:2500, Jackson ImmunoResearch). Western blots were imaged on ChemiDoc (Bio-Rad).

### Drug treatment

To perturb downstream signaling of p22, 8-9 hours post transfection, Latrunculin A (abcam), Wiskostatin (Sigma Aldrich), CK666 (Sigma Aldrich), smifH2 (CalBioChem), ZCL278 (Cayman Chemical), ML141 (Cayman Chemical) were added to complete media at specified concentrations. DMSO was used as vehicle control, and 36 hours post transfection the cells were imaged to quantify the extent of cell-cell fusion.

### Co-immunoprecipitation

Vero cells were transfected with specified plasmids. Vero cells harvested 24-36 hours post transfection. Cells were washed with PBS, and lysed with lysis buffer (150 mM NaCl, 25 mM HEPES pH 7.4, 1 mM EDTA, 0.5% NP-40, 1x HALT protease inhibitor (Thermo Fisher Scientific) for 30 min at 4°C, and scraped and bath sonicated in ice for 3 min. Cell debris was pelleted at 17,900 rcf for 10 min. Supernatant was incubated with 7.5 μl of washed GFP-Trap beads (Chromotek) overnight at 4°C. The beads were washed with lysis buffer five times, before boiled in Laemmli sample buffer and separated on 4-20% acrylamide gradient gels by SDS-PAGE. Proteins were transferred onto nitrocellulose membrane and probed with primary antibodies, α-ITSN (1:1000, Clone 29, BD Biosciences), α-GFP (1:5000, gp1544, Life Technologies), and secondary antibodies, α-mouse HRP (1:5000, Jackson Labs), α-rabbit HRP (1:5000, 65-6120, Thermo Fisher), α-rat AlexaFluor 647(1:5000, Life Technologies). Western blots were imaged on ChemiDoc (Bio-Rad).

### Protein purification

N-terminus GST-tagged SH3A, SH3B, SH3C, SH3D, SH3E domains from human intersectin was expressed in Rosetta for 4 hours at 37C. Cells were resuspended in lysis buffer (25 mM HEPES pH 7.2, 150 mM NaCl, 1 mM DTT, 1x PMSF) and lysed by sonication. Cell debris was pelleted and the supernatant was bound onto glutathione resin (GBiosciences). Resin was washed with 10 column volumes of wash buffer (25 mM HEPES pH 7.2, 150 mM NaCl, 1 mM DTT) and protein was eluted with elution buffer (25 mM HEPES pH 7.2, 150 mM NaCl, 1 mM DTT, 30mM glutathione). Protein was desalted into storage buffer (25 mM HEPES pH 7.2, 150 mM NaCl, 1 mM DTT, 10% glycerol) and flash frozen.

### SH3 binding assay

HEK293T cells were transfected with pcDNA3.1-p22-myc with TransIT-293 according to manufacturer’s instructions. 24 hours post transfection, the cells were washed with PBS and lysed by incubating in lysis buffer (150 mM NaCl, 25 mM HEPES pH 7.4, 1 mM EDTA, 0.5% NP-40, 1x HALT protease inhibitor (Thermo Fisher Scientific) for 30 min in 4°C and bath sonicated for 3 min. Cell debris was pelleted at 17,900 rcf for 15 min and the supernatant was bound to α-myc beads (Chromotek) for 4 hours at 4°C. Beads were washed twice wash buffer and (150 mM NaCl, 25 mM HEPES pH 7.2, 1 mM EDTA, 0.5% NP-40) and incubated with 30 μM of GST-SH3A, GST-SH3B, GST-SH3C, GST-SH3D, GST-SH3E for an hour at 4C. Beads were washed twice with (1 M NaCl, 25 mM HEPES pH 7.4, 0.1% NP-40, 1 mM EGTA, 5 mM MgCl_2_) and three times with (100 mM NaCl, 25 mM HEPES pH 7.4, 0.1% NP-40, 1 mM EGTA, 5 mM MgCl_2_). The beads were washed with lysis buffer five times, before boiled in Laemmli sample buffer and separated on 4-20% acrylamide gradient gels by SDS-PAGE. Proteins were transferred onto nitrocellulose membrane and probed with primary antibodies, α-GST (1:5000, abcam).

### Flow cytometry

Cells were transfected with p14-WT-GFP, p22-WT-GFP and chimera-GFP. 24 hours post transfection, the cells were lifted with short treatment of 0.05% trypsin, neutralized with full media and the GFP intensity was analyzed with Attune (Thermo Fisher). GFP-expressing population was identified by comparing with non-transfected cells, and the average GFP intensity of the GFP-expressing population was calculated using FlowJo.

## Acknowledgments

The authors would like to thank Fletcher Lab members, especially Andrew Harris, for useful feedback and technical consultation; Conklin Lab at UCSF for providing WTC10 hiPSC line; Lavis Lab at Janelia for providing JF635 HaloTag ligand; Dr. Sun Hae Hong for generating the AP2-tagRFP-T hiPSC line; the UC Berkeley QB3 MacroLab for purified S. pyogenes NLS-Cas9; Cancer Research Laboratory Flow Cytometry Facility for hiPSC sorting. This work is supported by R01GM114671 and R01GM134137 from NIGMS (D.A.F.), the Chan Zuckerberg Biohub (D.A.F.), DBI-1548297 from NSF (D.A.F.), and NIH MIRA R35GM118149 (D.G.D.). K.M.C.C. was funded by a NSF-GRFP fellowship. D.A.F. is a Chan Zuckerberg Biohub investigator. A.L.A. was supported by a NIH F31 grant (1F31GM128325-01). J.M. is supported through the NYU Margaret and Herman Sokol Fellowship, NYU Horizon Fellowship, and German Academic Fellowship Foundation (Studienstiftung) funding. J.M. and D.S. thanks the Boehringer Ingelheim Fonds for generously supporting his participation in MBL Physiology 2019. M.J. is supported by an American Heart Association postdoctoral fellowship (18POST34000029). D.S. is supported by a Boehringer Ingelheim Fonds PhD fellowship. S.S. was funded by a LSRF fellowship. The authors thank the Marine Biological Laboratory Physiology Course 2019 for providing an opportunity to explore initial ideas that formed the basis of this study, and we thank the Burroughs Wellcome Fund for supporting post-course fellowships for A.A. and J.M. to continue the project at UC Berkeley.

## Supplementary Figures

**Supplementary Figure 1.**
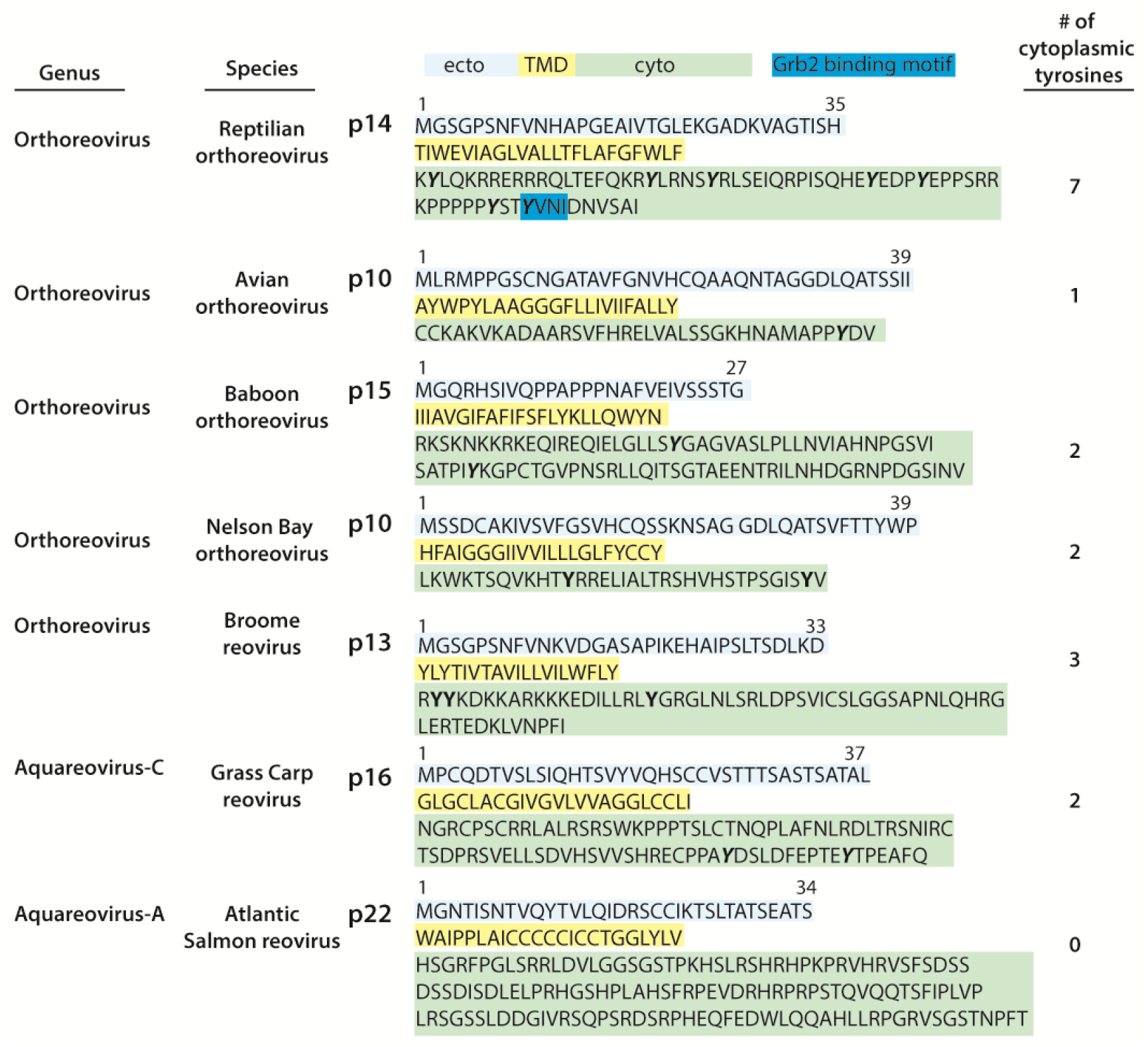
Amino acid sequence of FAST proteins. Amino acid sequence of p14 (Reptilian orthoreovirus, Q80FJ1), p10 (Avian reovirus-176, Q77ND6), p15 (Baboon orthoreovirus, Q918V6), p10 (Nelson Bay orthoreovirus, Q9J1B2), p13 (Broome virus, D6MM29), p16 (Grass Carp reovirus, Q8JU66) and p22 (Atlantic salmon reovirus-Canada 2009, C0L0N0) with known Grb2 binding motifs labeled in cyan and predicted topology and domains notated. The number of cytoplasmic tyrosines and virus genus, species/strain is shown.

**Supplementary Figure 2.**
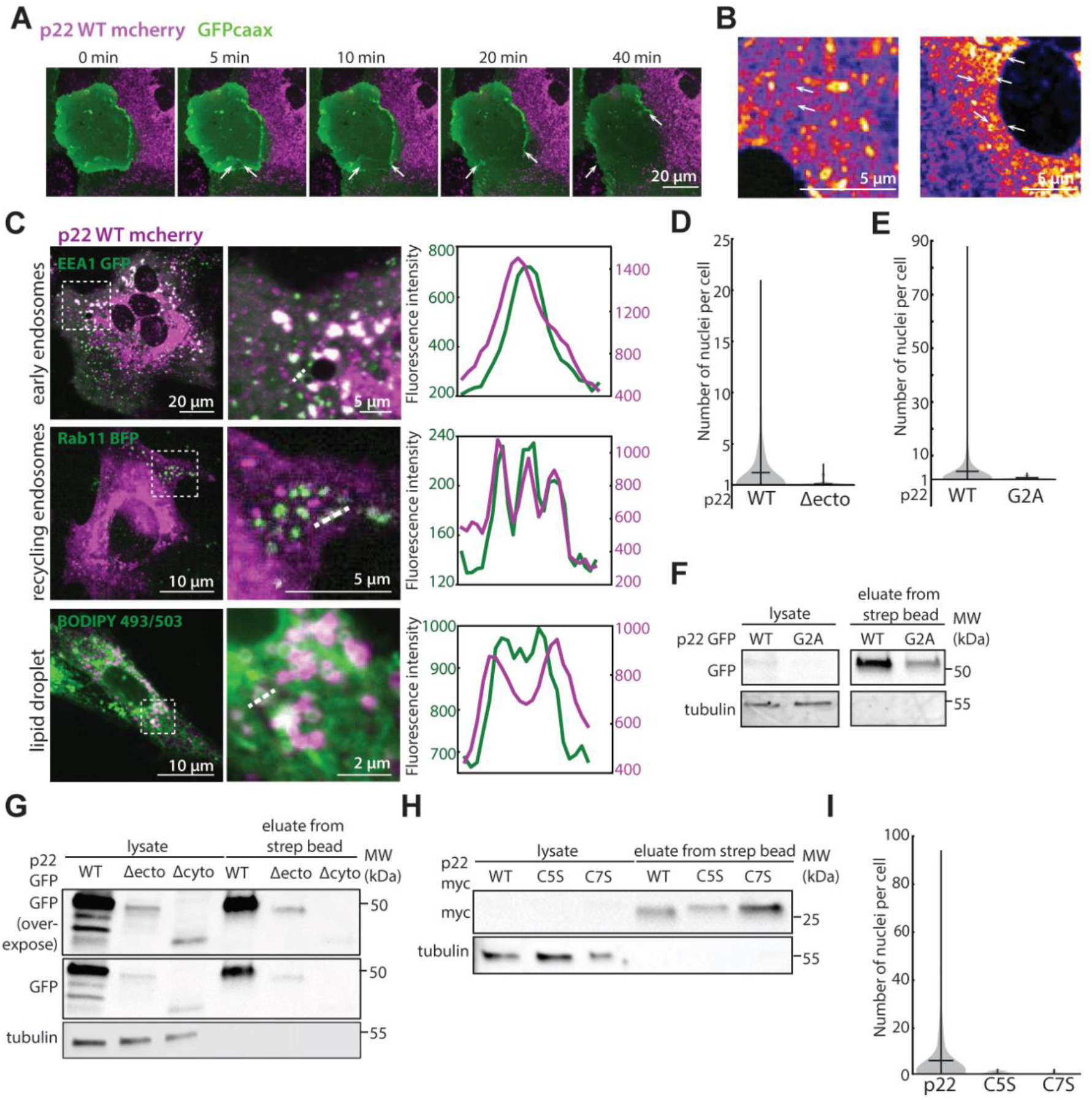
Intracellular localization of p22 and quantification of p22 transmembrane and ectodomain mutants. (A) Confocal image timeseries of p22-mcherry mCherry (magenta) expressing cell fusing with a neighboring naïve cell. Plasma membrane is visualized with GFP-caax (green) and pore expansion is denoted with white arrows (B) Regions from Figure 1D are magnified and vesicles are denoted with white arrows. (C) Representative confocal images of p22-mcherry (magenta) co-expressed with EEA1-GFP (green) and Rab11-BFP (green) to label early and recycling endosomes. Lipid droplets were labeled with BODIPY 493/503 (green). Regions boxed are magnified and fluorescence intensity of line scan of dotted line is shown. (D) Distribution and mean of number of nuclei in p22-WT and p22-G2A expressing cells from three independent transfections. (E) Distribution and mean of number of nuclei in p22-WT and p22-Δecto expressing cells from three independent transfections. (F) Western blot from surface biotinylation of GFP-tagged p22-WT and p22-G2A expressing cells. This is the same blot as in Supp. Figure 3E. (G) Western blot from surface biotinylation of GFP-tagged p22-WT, p22-Δecto and p22-Δcyto expressing cells. (H) Western blot from surface biotinylation of myc tagged p22-WT, p22 C5S and p22 C7S expressing cells. (I) Distribution and mean of number of nuclei in p22-WT, p22-C5S, and p22-C7S expressing cells from three independent transfections.

**Supplementary Figure 3.**
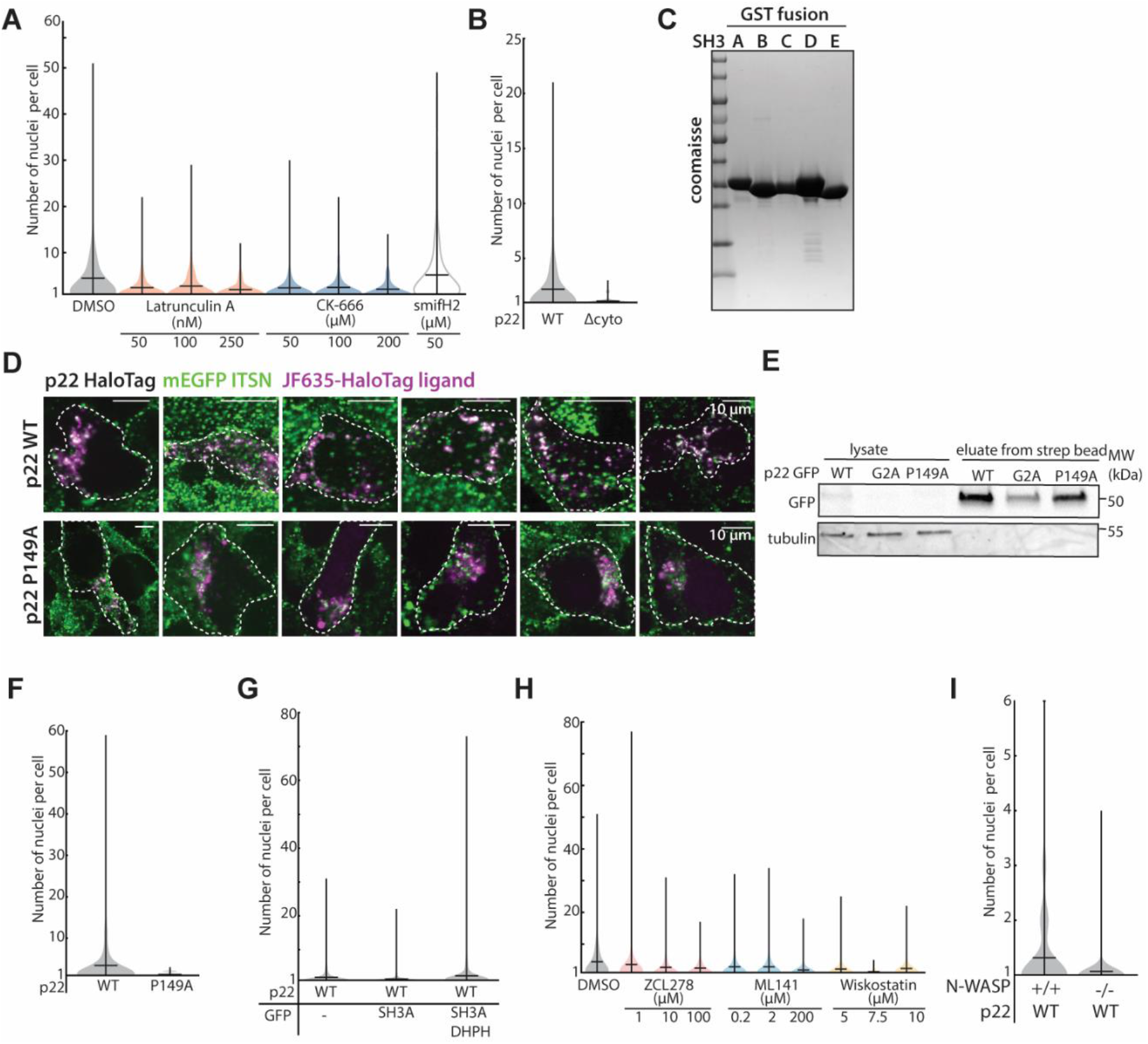
Quantification of p22 mutants and cytoskeletal drug treated p22-expressing cells. (A) Distribution and mean of number of nuclei in p22-WT expressing cells treated with Latrunculin A, CK-666, and smifH2 from three independent transfections. (B) Distribution and mean of number of nuclei in p22-WT and p22-Δcyto expressing cells from three independent transfections. (C) Coomassie stained acrylamide gel of purified GST-tagged SH3 domains of Intersectin-1. (D) Confocal images of endogenously mEGFP-tagged Intersectin-1 (green) cells expressing p22-WT-HaloTag (magenta) or p22-P149A-HaloTag (magenta) conjugated with JF635. The periphery of cells are outlined with white dotted line. (E) Western blot of surface biotinylation of GFP-tagged p22-WT, p22-G2A and p22-P149A. The p22-WT and p22-G2A lanes are reproduced in Supplementary Figure 2F. (F) Distribution and mean of number of nuclei in p22-WT and p22-P149A expressing cells from three independent transfections. (G) Distribution and mean of number of nuclei in p22-WT expressing cells co-expressing GFP alone or SH3A-GFP from 3 independent transfections. (H) Distribution and mean of number of nuclei in each p22-WT expressing cell treated with ZCL278, ML141, and Wiskostatin from three independent transfections. (I) Distribution and mean of number of nuclei in p22-WT expressing wild-type cells or N-WASP null cells from three independent transfections.

**Supplementary Figure 4.**
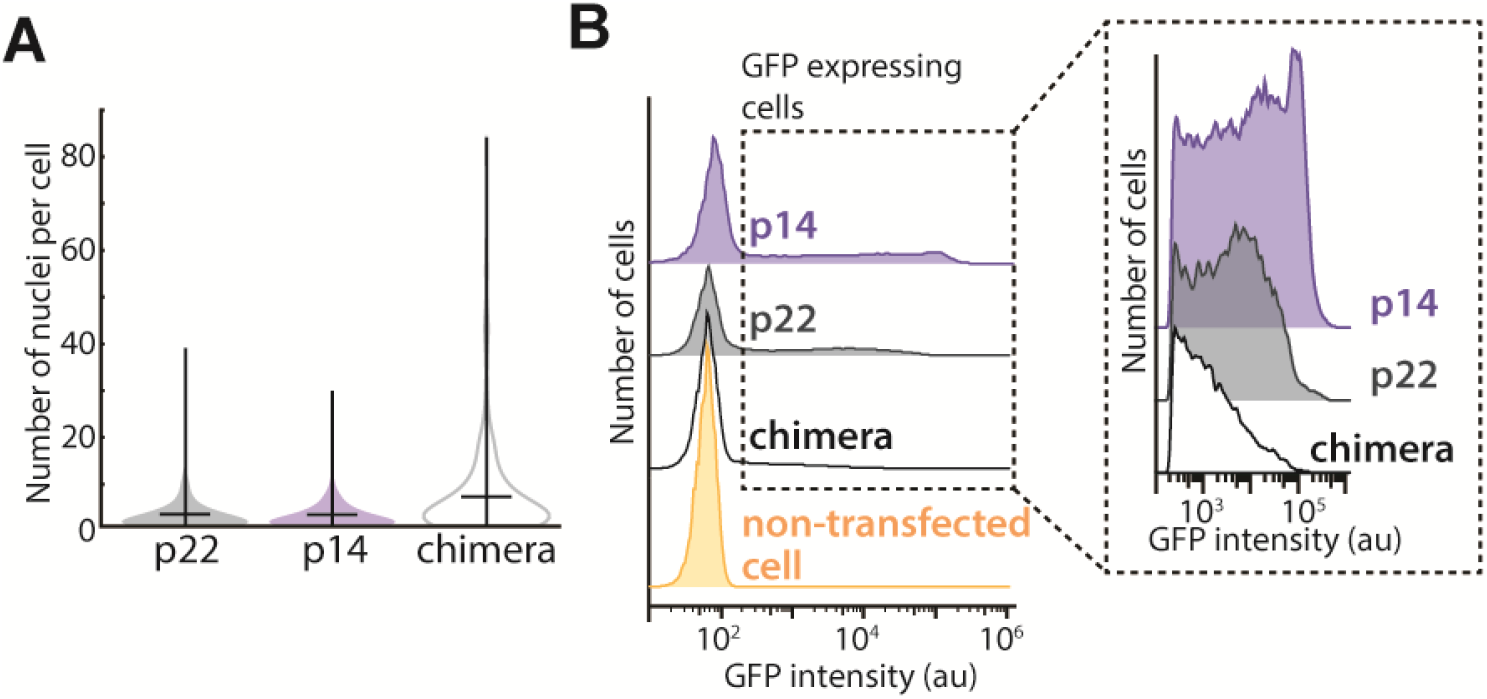
p14/p22 chimeric fusogen expression and distribution of number of nuclei in chimera-expressing cells. (A) Distribution of number of nuclei in p22, p14, and p14/p22 chimera-expressing cells from three independent transfections. (B) Distribution of GFP intensity of cells expressing GFP-tagged p14, p22 and p14/p22 chimera and non-transfected cells by flow cytometry. Boxed regions are magnified.

**Supplementary Figure 5.**
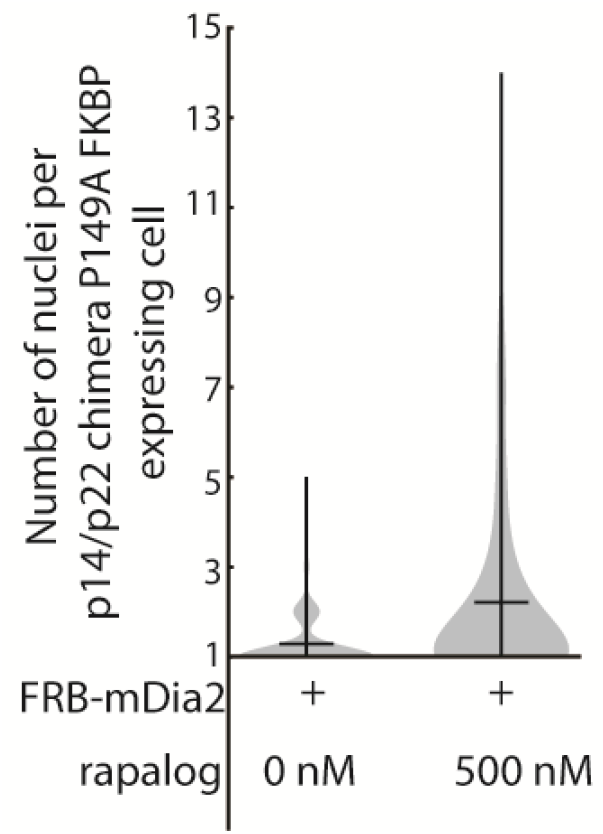
Quantification of fusion in chimera mutant coupled to a formin. Distribution and mean of number of nuclei in p14/p22 chimera-P149A-FKBP and FRB-mDia2 expressing cell treated with rapalog from three independent transfections.

**Supplementary Figure 6.**
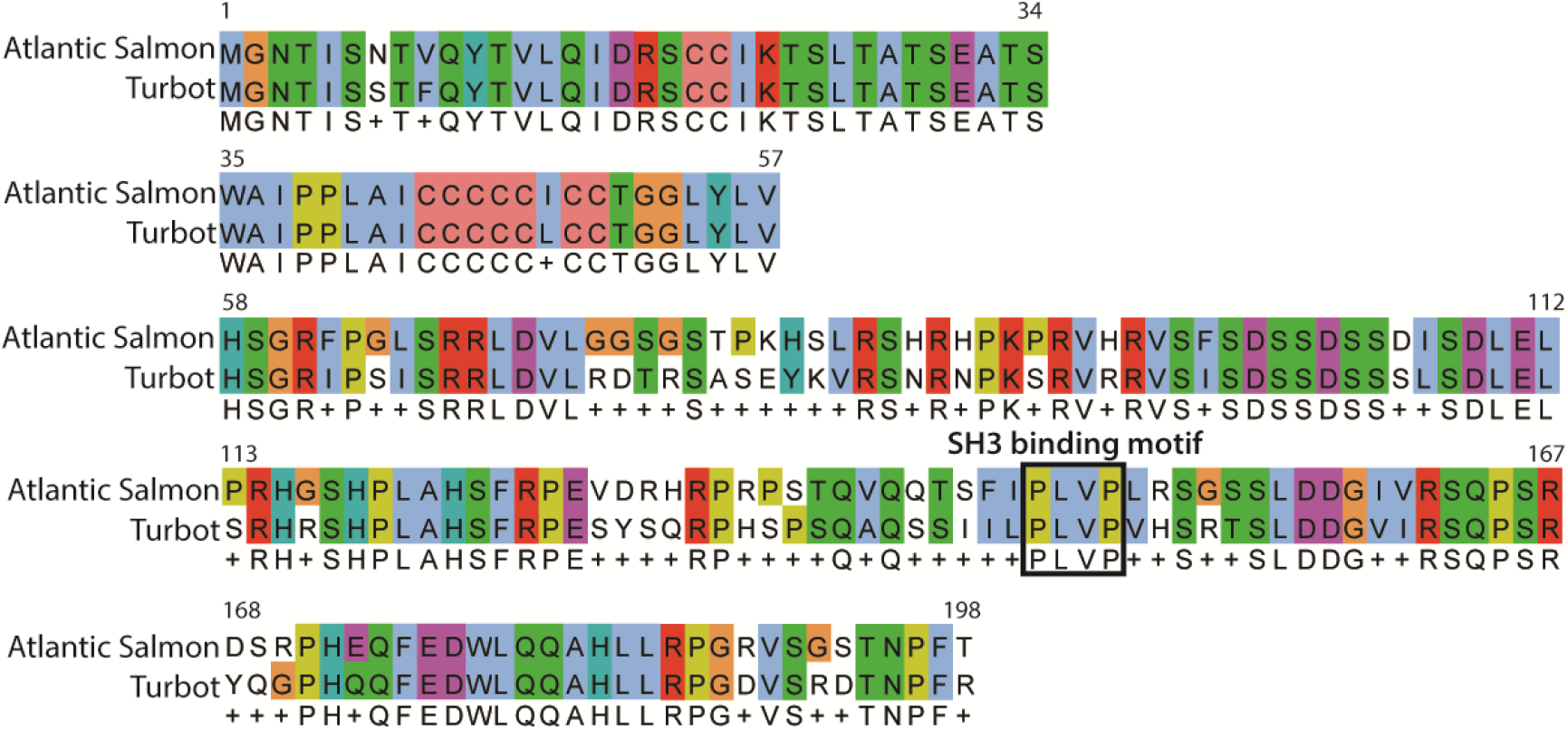
Sequence alignment of FAST proteins from Turbot reovirus and Atlantic salmon reovirus using ClustalW. Sequence alignment of FAST proteins from Turbot reovirus (ADZ31982.1) and Atlantic salmon reovirus (C0L0N0) using ClustalW.

## Supplementary Videos

**Video 1.** Confocal timelapse of a Vero cell expressing p22-mcherry (magenta) and GFPcaax (green) fusing with a Vero cell expressing only GFPcaax (green). Scale bar is 20 μm.

**Video 2.** Confocal timelapse of a Vero cell expressing chimera-P149A-mCherry-FKBP (magenta), FRB-ΔGBD-mDia2 and Lifeact-GFP (green) upon addition of 500 nM rapalog. Scale bar is 10 μm.

**Video 3.** Confocal timelapse of a magnified region of Video 2. Vero cell expressing chimera-P149A-mCherry-FKBP (magenta), FRB-ΔGBD-mDia2 and Lifeact-GFP (green) upon addition of 500 nM rapalog. Scale bar is 10 μm.

